# Selective conservation of symbiont cell-surface glycans across generations in a vertically transmitting coral

**DOI:** 10.64898/2026.04.21.719984

**Authors:** Giada Tortorelli, Nerissa L. Fisher, Alyssa C. Varela, Sabrina L Rosset, Immy A. Ashley, Eva Majerová, Khalil Smith, Kira Hughes, Crawford Drury

## Abstract

Coral resilience under climate change depends on the stability of coral–Symbiodiniaceae symbioses. While vertically transmitting corals inherit symbionts directly from parental colonies, the extent to which symbiont cellular traits are conserved across life stages remains unclear. Here, we examined cell-surface glycan profiles of Symbiodiniaceae in parental colonies and eggs of the coral *Montipora capitata*. Glycan signatures were structured by symbiont genus and differed between life stages, with mannose/glucose- and galactose-containing glycoproteins as primary drivers of variation. Despite life-stage differences, parent–offspring comparisons revealed significant conservation of glycan profiles, indicating intergenerational transmission of symbiont cellular traits that differed between *Cladocopium* and *Durusdinium* and were driven by distinct glycan classes. These results suggest that vertical transmission preserves key recognition-relevant glycans while allowing flexibility in other symbionts’ surface traits, providing a mechanistic basis for symbiosis stability.

## Introduction

Reef-building corals form an obligate endosymbiosis with dinoflagellates of the family Symbiodiniaceae, a partnership that underpins the productivity and persistence of coral reef ecosystems (1). Symbiodiniaceae provide photosynthetically derived carbon to the host, supporting metabolism, calcification, and reef growth, while receiving inorganic nutrients and a protected intracellular niche. This metabolic coupling represents one of the most ecologically successful symbioses in the ocean and has persisted for hundreds of millions of years, shaping the evolution and biogeography of coral reefs (2,3).

Despite this long evolutionary history, the coral–Symbiodiniaceae symbiosis is highly sensitive to environmental change. Elevated sea surface temperatures destabilize this association, causing bleaching through impaired photosynthesis, nutritional decoupling (4), and reactive oxygen–mediated symbiont loss (5,6). Recurrent marine heatwaves have increased the frequency and severity of mass bleaching, driving coral mortality, declines in coral cover, and community shifts. As a result, a substantial proportion of reefs have already been degraded, with further losses projected under continued warming (7).

Considering Symbiodiniaceae are indispensable to coral fitness, the mode by which corals acquire their symbionts is a key determinant of coral biology. Symbiont transmission occurs along a continuum between horizontal acquisition from the environment and vertical inheritance from the parent colony, with important consequences for host–symbiont specificity, flexibility, and resilience (8). Horizontally transmitting corals acquire symbionts from the surrounding seawater, enabling uptake of locally adapted partners with reduced fidelity. In contrast, vertically transmitting corals provision symbionts via the eggs, ensuring early symbiosis and high intergenerational fidelity (9), though potentially limiting rapid adjustment to environmental change. These contrasting transmission strategies are therefore expected to influence not only the composition of symbiont communities but also holobiont performance under stress (10).

The reef-building coral *Montipora capitata* is a dominant species in Kāneʻohe Bay, Hawaiʻi, and a key model for coral–Symbiodiniaceae symbiosis under environmental stress. It is a broadcast spawner with vertical transmission, releasing egg–sperm bundles that develop into larvae containing maternally derived symbionts (10,11). This strategy establishes symbiosis early and links parental and offspring communities, yet variation across colonies and environments indicates both inherited and environmental influences on symbiont dynamics.

Across its range, *M. capitata* associates primarily with symbionts from the genera *Cladocopium* and *Durusdinium*, which differ in physiological performance and stress tolerance (12,13). *Cladocopium* is generally considered more thermally sensitive but can support higher rates of photosynthesis and growth under stable environmental conditions (2,12,14,15). In contrast, *Durusdinium* is widely recognized for its enhanced thermal tolerance and resistance to bleaching, although this resilience is often associated with trade-offs in host growth or metabolic efficiency (2,12,14–16). As a result, the relative abundance of these symbionts can influence coral performance across environmental gradients, particularly under elevated temperature regimes (14,15), providing a framework to investigate how symbiont traits are maintained across life stages.

At the cellular level, the establishment and maintenance of the coral–Symbiodiniaceae symbiosis is mediated by molecular recognition processes often described as a “lock-and-key” mechanism, in which symbiont cell-surface glycans interact with host lectins to enable selective recognition, uptake, and retention of compatible symbiotic partners (1,17). Experimental evidence has shown that lectin–glycan interactions influence symbiont acquisition, with certain carbohydrate motifs promoting or inhibiting symbiont colonization, thereby contributing to host–symbiont specificity (14,18–28). Most of our current understanding of these recognition mechanisms comes from horizontally transmitting systems, where symbiont uptake from the environment requires active discrimination among diverse potential partners and lectin–glycan interactions mediate partner selection at the host–cell interface. How these molecular recognition processes operate in vertically transmitting corals remains poorly understood. Because symbionts are inherited directly from the parent colony via symbiont-provisioned eggs, the need for de novo partner acquisition is assumed to be reduced. However, recognition mechanisms may still be essential for regulating symbiont retention, proliferation, and quality control across life stages. In this context, lectin–glycan interactions may function not only in initial symbiont recognition but also in maintaining symbiotic fidelity and mediating intergenerational continuity of host–symbiont associations.

In *M. capitata* from Kāneʻohe Bay, cell-surface glycan profiles of *Cladocopium* and *Durusdinium* differed between genera and were significantly altered by temperature and oxidative stress, suggesting that deviations from baseline glycan signatures may contribute to the breakdown of specific partnerships and the plasticity of symbiosis under stress (14).

Here, we investigate how symbiont cell-surface glycan profiles are maintained and structured across life stages in a vertically transmitting coral. Using *M. capitata* as a model system, we test whether lectin-binding signatures of Symbiodiniaceae are conserved between parental colonies and their offspring, and whether this signature varies between *Cladocopium* and *Durusdinium*, providing insight into the role of lectin–glycan mechanisms in maintaining symbiosis across generations.

## Materials and methods

### Coral collection, Symbiodiniaceae isolation, and qPCR

*M. capitata* colonies in Kāneʻohe Bay, Oʻahu, were selected based on visual health and colony size to maximize the likelihood of spawning. Colonies were tagged *in situ*, and thumb-sized fragments (*N* = 3-5) were collected from each colony for symbiont isolation and molecular analyses.

To characterize the algal symbiont community, fragments were sampled using a 5 mm diameter dermal curette, immediately snap-frozen in liquid nitrogen, and stored at −80°C until processing. Genomic DNA was extracted using the Zymo Quick-DNA kit (Fisher Scientific, cat. no. 50-444-148) following the manufacturer’s protocol. Quantitative PCR (qPCR) assays targeting clade-specific actin loci were used to quantify the relative abundance of *Cladocopium* and *Durusdinium* in corals (12,29). All qPCR reactions were performed on a StepOnePlus Real-Time PCR System (Applied Biosystems). Reactions were carried out in 10 µL volumes containing 5 µL TaqMan Genotyping Master Mix and 1 µL genomic DNA template. The *M. capitata* assay contained 1 µM forward primer (PaxC-F, 5ʹ-GTGCAGGTGAGATTGAGTCTTATAACA-3ʹ), 1.5 µM reverse primer (PaxC-R, 5ʹ- CGGTTGAGCTTCGCTAAACAG-3ʹ), and 2 µM TaqMan probe (PaxC-Probe, 5ʹ-FAM-CAGTTCTTCCAACAATG-MGB-3ʹ). The multiplexed *Cladocopium*/*Durusdinium* assay contained 1 µM *Cladocopium* forward primer (CActF, 5ʹ-CCAGGTGCGATGTCGATATTC-3ʹ), 1.5 µM *Cladocopium* reverse primer (CActR, 5ʹ-TGGTCATTCGCTCACCAATG-3ʹ), and 2 µM *Cladocopium* probe (CActProbe, 5ʹ-VIC-AGGATCTCTATGCCAACG-MGB-3ʹ), together with 1 µM *Durusdinium* forward primer (DActF, 5ʹ-GGCATGGGGTAAGCACTTCTT-3ʹ), 1.5 µM *Durusdinium* reverse primer (DActR, 5ʹ-GATCCTTGAACTAGCCTTGGAAAC-3ʹ), and 2 µM *Durusdinium* probe (DActProbe, 5ʹ-6FAM-CAAGAACGATACCGCC-MGB-3ʹ). All samples were run in triplicate for each assay over 40 cycles. Thermal cycling conditions consisted of an initial incubation at 50°C for 2 min and 95°C for 10 min, followed by 40 cycles of denaturation at 95°C for 10 s and annealing/extension at 60°C for 1 min. Cycle threshold (CT) values were determined using the StepOnePlus software with a baseline interval of cycles 6–23 and a fluorescence threshold of Rn = 0.01. Positive amplification was scored only when technical replicates yielded CT values < 40 and no amplification was detected in no-template controls. CT values were corrected for differences in fluorescence intensity among the three reporter dyes associated with the TaqMan probes. Symbiont-to-host (S/H) cell ratios for *Cladocopium* and *Durusdinium* were calculated as 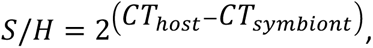 and values were normalized for differences in probe fluorescence intensity and target locus gene copy number. Total S/H ratios were calculated as the sum of *Cladocopium* and *Durusdinium* S/H ratios (12,29).

In total, 16 *M. capitata* colonies were selected, including 6 dominated by *Cladocopium* (genotypes ID 162, 176, 177, 190, 214, 236) and 10 dominated by *Durusdinium* (169, 185, 197, 203, 208, 228, 231, 243, 249, 672). Colonies were used for downstream analyses if their symbiont community was composed of > 99.5% of a single genus; composition data can be found in Supplementary Table 1.

### Adult samples preparation

Algal symbionts were freshly isolated from a subset of 11 parental colonies (*Cladocopium* genotypes 162, 177, 190, 214, 236, and *Durusdinium* genotypes 185, 203, 208, 228, 231, 243) by airbrushing tissue in 0.2 μm filtered seawater (FSW). Logistical constraints prevented us from sampling all parental colonies. The resulting homogenate was dissociated using a sterile 22-gauge needle and syringe to obtain single cells, centrifuged at 1,000 × g for 5 min to pellet symbiont cells and resuspended in FSW. This wash step was repeated five times to remove residual host material. Symbiont cell pellets were fixed in 2% paraformaldehyde in 1x PBS and stored in the dark at 4°C until further processing.

### Spawning and egg samples preparation

Tagged *M. capitata* colonies were transported to the Hawaiʻi Institute of Marine Biology (HIMB) in Kāneʻohe Bay three days prior to the new moon in May and June 2025 and maintained in ambient flow-through seawater tanks under natural light conditions. Gamete bundles were collected from individual spawning coral colonies using a net on May 28 and June 26, 2025. Buoyant gamete bundles were gathered in individual 50mL conical tubes and were gently agitated to dissociate bundles and release eggs and sperm. Eggs were repeatedly washed with FSW to remove residual sperm and debris, then aliquoted into 1.5 mL microcentrifuge tubes.

To isolate algal symbionts from the eggs, samples were vortexed horizontally at maximum speed for 5 min and centrifuged at 1,000 × g for 5 min. The supernatant was discarded, and the symbiont pellet was resuspended in FSW using a sterile 22-gauge needle and syringe. This wash cycle was repeated five times to discard eggs debris. Symbiont cell pellets from replicate samples were pooled, fixed in 2% paraformaldehyde in PBS, and stored in the dark at 4°C until further processing.

### Lectin staining and flow cytometry of Symbiodiniaceae cell-surface glycans

Fluorescent lectins were used to characterize glycans present on the cell-surface of symbiotic algae. The following lectins, purchased from Thermo Fisher Scientific, were used: ConA (concanavalin A; specific for D-mannose and D-glucose; cat. no. C21401), LTL (*Lotus tetragonolobus* lectin; specific for L-fucose; cat. no. L32480), PNA (peanut agglutinin; specific for D-galactose; cat. no. L21409), WGA (wheat germ agglutinin; specific for N-acetylglucosamine and N-acetylneuraminic acid; cat. no. W11261), PHAL (phytohemagglutinin-L from *Phaseolus vulgaris*; specific for N-acetylglucosamine β(1–2) mannopyranosyl residues; cat. no. L11270), and GSIB4 (isolectin B4 from *Griffonia simplicifolia*; specific for N-acetyl-D-galactosamine and α-D-galactosyl residues; cat. no. I21411). Lectins were conjugated with either Alexa Fluor 488 (ConA, PNA, WGA, PHAL, GSIB4) or FITC (LTL). Lectin-binding profiles of Symbiodiniaceae cell-surface glycans were quantified for symbionts isolated from the eggs of 6 *Cladocopium*-dominated and 10 *Durusdinium*-dominated colonies, and from adult tissue of 5 *Cladocopium*-dominated and 6 *Durusdinium*-dominated parental colonies.

Algal cell density was determined from quadruplicate counts using a Countess 3 FL Automated Cell Counter (Thermo Fisher Scientific). For each sample, 2.5 × 10⁵ fixed symbiont cells were incubated separately with each fluorescent lectin at a final concentration of 1 mg ml⁻¹ for 1 hour at room temperature in the dark. Three stained replicates were prepared for each biological replicate. Following incubation, cells were washed with 1× PBS by centrifugation at 3,000 × g for 5 min and resuspended in a final volume of 500 µl PBS (14).

Samples were processed on a CytoFlex S flow cytometer (Beckman-Coulter) for at least 90 s at a flow rate of 20 µl min^-1^ and data were analyzed using FloJo software (v10.10.0, BD BioSciences). Gain parameters were established before running experimental samples using unstained and stained algal cells. Algal cell populations were first gated by chlorophyll autofluorescence and Alexa Fluor 488 fluorescence (on the SYBR Green channel) based on excitation at 488 nm and detected by the PerCP channel (690/50 bandpass filter) and FITC channel (525/40 bandpass filter), respectively. The median fluorescence intensity (MFI) of the lectin-binding height signal (SYBR Green) was quantified using the gated algal population (Supplementary Fig. 1). Background fluorescence was corrected by subtracting the MFI of unstained cells from lectin-stained samples.

### Statistical analysis

All statistical analyses were conducted in R (v4.1.2). Coral colonies were classified according to their dominant symbiont genus based on qPCR estimates of *Cladocopium* and *Durusdinium* relative abundance following Cunning et al. 2013 (29).

Lectin-binding profiles of symbiont cell-surface glycans were quantified as median fluorescence intensity (MFI) for each of the six lectins (ConA, LTL, PNA, WGA, PHAL, GSIB4) and unstained controls. Lectin MFI values were corrected by subtracting the chlorophyll autofluorescence signal measured for each sample from the corresponding unstained negative control, thereby accounting for background fluorescence and isolating the lectin-specific binding signal. Multivariate differences in lectin-binding profiles between symbiont genera and sample origin (eggs vs. parents) were assessed using permutational multivariate analysis of variance (PERMANOVA; 999 permutations) and homogeneity of multivariate dispersion (β-dispersion) implemented in the *vegan* package (30). We used partial least squares discriminant analysis (PLS-DA) to evaluate the ability of each glycan’s z-scored MFI to distinguish symbiont types in eggs and parents using the *mixOmics* package (31).

To test the effects of lectin type and symbiont genus on lectin-binding profiles of symbionts isolated from eggs and parental coral colonies, we fitted a generalized linear model with a Student’s t error distribution using the *glmmTMB* package (32). The model included lectin type, symbiont genus, and host genotype nested within symbiont genus as fixed effects (MFI ∼ lectin × symbiont + symbiont/genotype). Model diagnostics were performed using the *DHARMa* package (33). Significance of fixed effects was assessed using analysis of variance (ANOVA), and *post hoc* pairwise comparisons were conducted using Tukey-adjusted contrasts implemented in the *emmeans* package (34).

Associations between lectin-binding profiles of symbionts isolated from eggs and parental colonies were examined using pairwise correlation analyses. Correlation matrices were constructed from z-scaled MFI values across all six lectins, and similarity between matrices was assessed using Mantel tests (999 permutations) implemented in the *vegan* package (30). Comparisons were performed between symbiont genera from eggs and between eggs and parents within each symbiont genus.

We compared Mahalanobis distances between matched parent–offspring pairs (POP) and unmatched comparisons (other) within each symbiont genus to assess whether lectin-binding profiles of symbiont cell-surface glycans were conserved between parental colonies and their offspring. Distances were analyzed using a generalized linear model including symbiont genus, comparison type (POP or other) and their interaction (distance ∼ symbiont × comparison type). Significance was assessed using analysis of variance (ANOVA), and *post hoc* pairwise comparisons were conducted using Tukey-adjusted contrasts implemented in the *emmeans* package (34). To assess transmission of each glycan, we used z-scaled MFI data as input for parent-offspring linear regressions separately for each lectin and symbiont type (egg ∼ parent), and an ANCOVA to assess differences in slope between symbiont types for each lectin.

## Results

### Symbiont genera and life stage differences in lectin-binding profiles

Multivariate analysis revealed clear separation in lectin-binding profiles of symbiont cell-surface glycans between symbiont genera at both life history stages. Principal component analysis showed distinct clustering of *Cladocopium* and *Durusdinium* in eggs (PERMANOVA, *P* = 0.001; Fig. 1A) and parents (PERMANOVA, *P* = 0.001; Fig. 1B), indicating genus-specific lectin-binding signatures. The lectin ConA contributed strongly to the separation between symbiont genera in both eggs and adults. WGA and PNA also strongly separated symbiont genera in eggs, but not in parental corals, suggesting a more complex glycan landscape in gametes than adults. PC2 captured substantial additional variation in eggs and within *Cladocopium* in adults. Multivariate dispersion was not different between symbiont genera in eggs (β-dispersion, *P* > 0.05; Fig. 1A) but was significantly higher in *Cladocopium-*hosting parents (β-dispersion, *P* = 0.043; Fig. 1B).

**Fig. 1.**
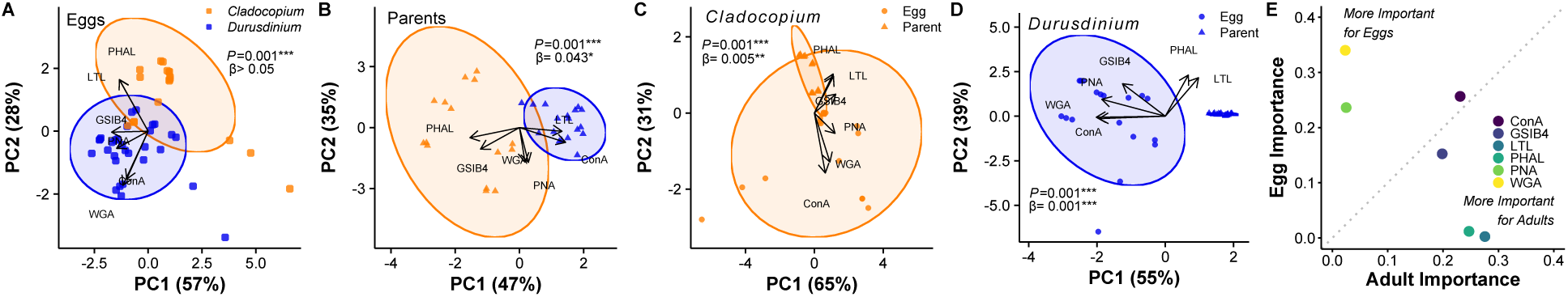
Lectin-binding profiles distinguish symbiont genera and host life stages. **A.** Principal component analysis (PCA) of lectin-binding intensities in *Cladocopium* (orange, *N* = 18) and *Durusdinium* (blue, *N* = 30) symbionts isolated from coral eggs. **B.** PCA of lectin-binding intensities in *Cladocopium* and *Durusdinium* symbionts isolated from parental colonies (*N* = 15 and *N* = 18, respectively). **C.** PCA of lectin-binding intensities in *Cladocopium* symbionts isolated from coral eggs (circles) and parental colonies (triangles) pairs (*N* = 15). **D.** PCA of lectin-binding intensities in *Durusdinium* symbionts isolated from coral eggs and parental colony pairs (*N* = 18). In A-D, vectors represent the contribution of individual lectins to the ordination. Ellipses represent the 95% confidence intervals around group centroids. Group separations were statistically significant (PERMANOVA *P* = 0.001). Asterisks denote adjusted *P* values (\**P*< 0.05, ** *P*< 0.01, *** *P*< 0.001). **E.** Partial least squares discriminant analysis (PLS-DA) of each lectin-binding intensity profile for symbionts in eggs and parents. Lectins: ConA (concanavalin A, specific for D-mannose and D-glucose), LTL (*Lotus tetragonolobus* lectin, specific for L-fucose), PNA (*Arachis hypogaea* lectin, specific for D-galactose), WGA (wheat germ agglutinin, specific for N-acetylglucosamine and N-acetylneuraminic acid), PHA-L (phytohemagglutinin-L from *Phaseolus vulgaris*, specific for N-acetylglucosamine β(1-2) mannopyranosyl) and GS-IB4 (isolectin from *Griffonia simplicifolia*, specific for N-acetyl-D-galactosamine and a-D-galactosyl residues).

Differences between symbionts isolated from eggs and parental colonies were detected in both genera (PERMANOVA, *P* = 0.001; Fig. 1C, D), indicating that variability in lectin-binding profiles also differed based on symbiont community. In *Cladocopium,* the separation was less pronounced, and confidence ellipses demonstrated some overlap (Fig. 1C), but in *Durusdinium,* egg- and parent-derived symbionts formed discrete clusters (Fig. 1D). Symbionts isolated from eggs had higher multivariate dispersion than symbionts isolated from parents in both genera (β-dispersion, *P* < 0.001; Fig. 1C, D). Partial least squares discriminant analysis (PLS-DA) showed that the lectins PNA and WGA were disproportionately important for distinguishing symbiont communities in eggs, while LTL and PHAL were more important in adults (Fig. 1E). The lectins ConA and GSIB4 were approximately equally important for distinguishing the symbiont community in eggs and parents.

Analysis of proportional median fluorescent intensity (MFI) of lectin-binding signal composition revealed that both symbiont genera from eggs and parental colonies were dominated by ConA and WGA signals, with relatively minor contributions from other lectins (Supplementary Fig. 2). In *Cladocopium* from eggs, genotype-level profiles were relatively consistent, with ConA contributing the majority of total signal across all genotypes (Supplementary Fig. 2C). *Durusdinium* from eggs exhibited greater variability among genotypes, particularly in the relative contribution of WGA and secondary lectins (Supplementary Fig. 2D), suggesting increased heterogeneity in lectin-binding profiles within this genus.

### Lectin-specific differences in symbiont cell-surface glycan binding

To identify lectin-specific differences underlying multivariate patterns, we compared median fluorescence intensity (MFI) across symbiont genera using generalized linear models. In eggs and parental colonies, lectin type, symbiont genus, and their interaction, all significantly influenced MFI (GLM, *P* < 0.0001 for all terms), indicating that glycan-binding differences are lectin-dependent and vary between symbiont genera (Fig. 2). Host genotype nested within symbiont genus also contributed significantly to variation in MFI (GLM, *P* < 0.001). Consistent with these effects, *Durusdinium* exhibited higher MFI than *Cladocopium* for several lectins (Fig. 2, Supplementary Fig. 2), with the strongest differences observed in eggs in ConA and WGA (*P* < 0.0001), as well as GSIB4 (*P* = 0.003) and PNA (*P* = 0.012), and in parents in ConA (*P* = 0.012), GSIB4 (*P* = 0.045) and PHAL (*P* = 0.042).

**Fig. 2.**
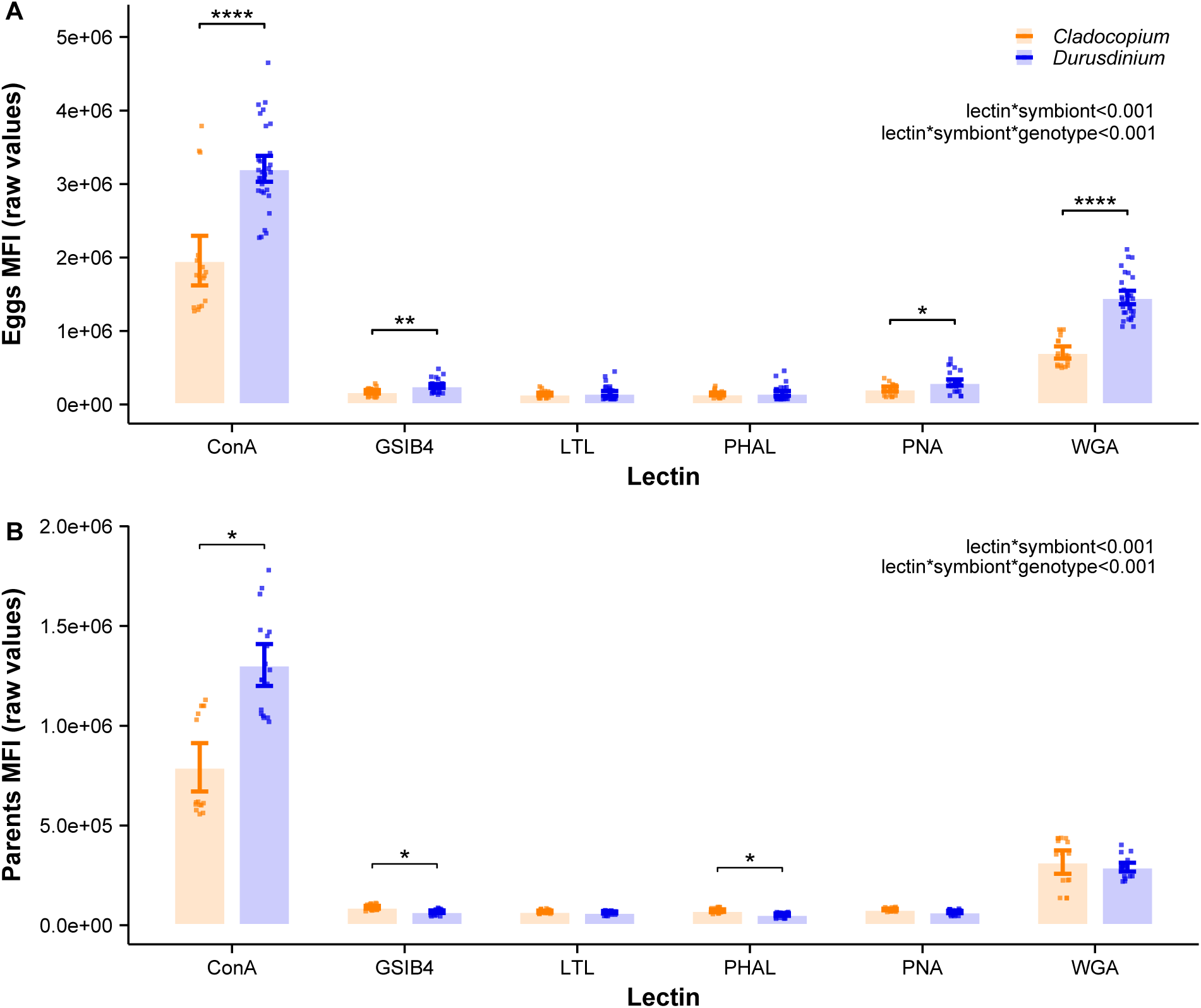
Lectin-binding intensity differs between genera in symbionts isolated from coral eggs and parental colonies. **A.** Median fluorescence intensity (MFI) raw data of lectin-binding measured in symbiont cells from the genera *Cladocopium* (orange, *N* = 18) and *Durusdinium* (blue, *N* = 30) isolated from coral eggs. **B.** Median fluorescence intensity (MFI) raw data of lectin-binding measured in symbiont cells from the genera *Cladocopium* (*N* = 15) and *Durusdinium* (*N* = 18) isolated from parental coral colonies. Points represent genotype-level mean values. Bars indicate the mean across genotypes, and error bars represent the standard error (SE). Asterisks denote adjusted *P* values (\**P*< 0.05, ** *P*< 0.01, *** *P*< 0.001).

### Correlation structure of lectin-binding profiles across symbiont genera and life stages

To characterize relationships between lectin-binding profiles of symbiont cell-surface glycans, we constructed correlation matrices for each symbiont genus and coral life stage (Fig. 3). In *Cladocopium*, correlations were exclusively positive in eggs and parents, although some lectin pairs showed weak or no association. The overall correlation structure was significantly different between life history stages in *Cladocopium* (Mantel *r* = 0.861, *P* = 0.001; Fig. 3A). In contrast, *Durusdinium* exhibited more variable correlation patterns, with both positive and negative associations observed in egg- and parent-derived symbionts. Generally weaker correlations within PNA and ConA in both eggs and parents with *Durusdinium* contributed to a lower correlation between sample types, although the overall correlation structure was still significantly different between life history stages (Mantel *r* = 0.505, *P* = 0.044; Fig. 3B). Correlation matrices were significantly different between symbiont genera in samples from parents (Mantel *r* = 0.726, *P* = 0.003) and eggs (Mantel *r* = 0.777, *P* = 0.019).

**Fig. 3.**
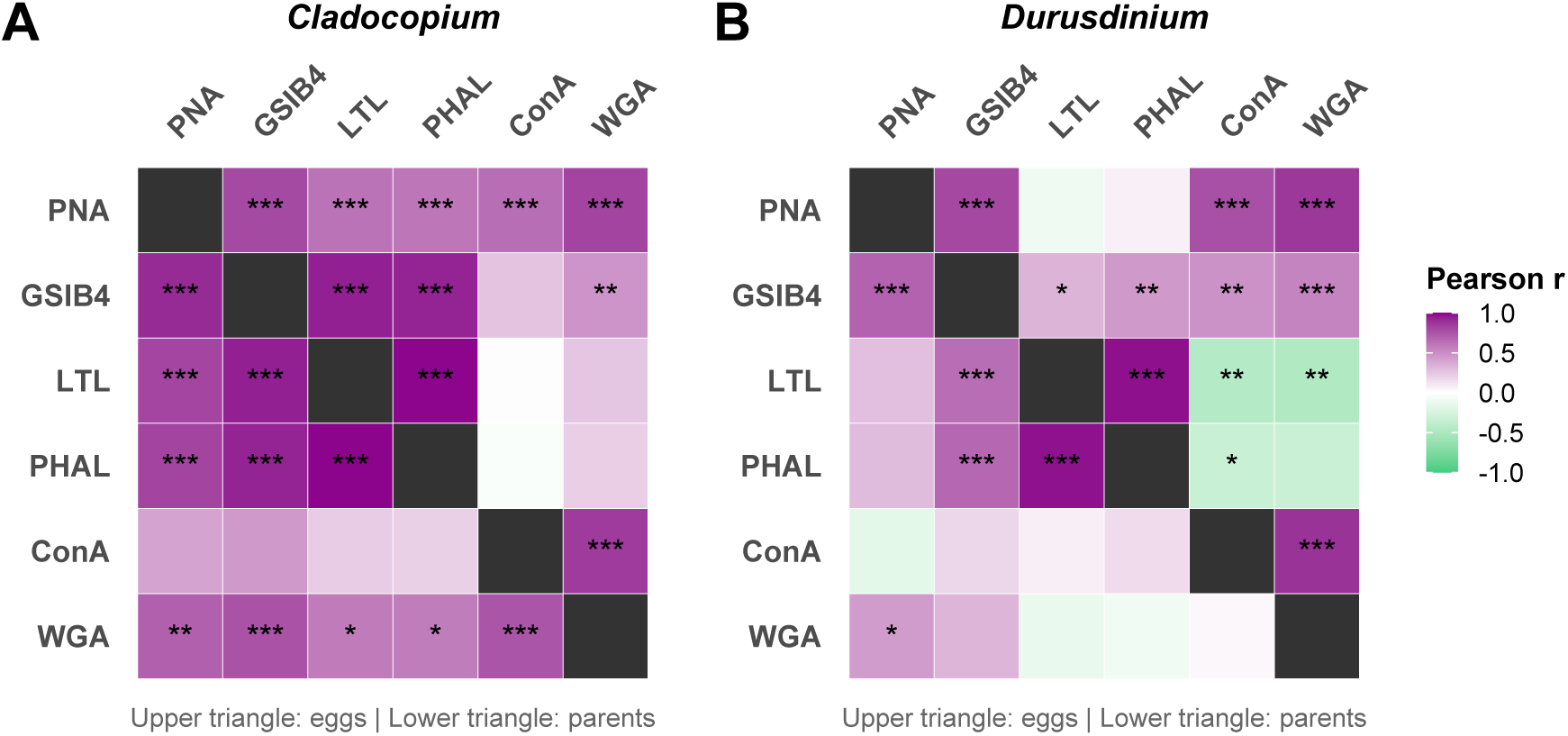
Pairwise Pearson correlations between lectin-binding intensities differ between symbiont genera and host life stages. Heatmaps show pairwise Pearson correlations in lectin-binding intensities measured in **A.** *Cladocopium* and **B.** *Durusdinium* cells isolated from coral eggs (upper triangles) and parental colonies (bottom triangles). Color scale represents the strength and direction of correlations (purple: positive, green: negative). Similarity between lectin correlation matrices was assessed using Mantel tests based on 999 permutations (*P_C-eggs vs C-parents_*= 0.001; *P_D-eggs vs D-parents_*= 0.044; *P_C-eggs vs D-eggs_*= 0.003; *P_C-parents vs D-parents_*= 0.019). Asterisks within cells indicate significant correlation coefficients between lectin-binding intensities (\**P*< 0.05, ** *P*< 0.01, *** *P*< 0.001).

### Host genotype-specific similarity in symbionts lectin-binding profiles

To assess whether lectin-binding profiles of symbiont cell-surface glycans were conserved between parental colonies and their offspring, we compared Mahalanobis distances between matched parent–offspring pairs (POP) and unmatched comparisons (other) within each symbiont genus. Distances were analyzed using a generalized linear model including symbiont genus, comparison type (POP or other), and their interaction. Distance was significantly influenced by the interaction between symbiont genus and comparison type (*χ*² = 23.26, *df* = 1, *P* < 0.001), but the main effect of symbiont and comparison type alone were not significant (Symbiont *χ*² = 0.07, *df* = 1, *P* = 0.795; Group *χ*² = 1.61, *df* = 1, *P* = 0.205). *Post hoc* comparisons revealed that distances were significantly smaller in POP than in unrelated pairs in *Cladocopium* (*P* = 0.003) and significantly higher than in unrelated pairs in *Durusdinium* (*P* < 0.0001; Fig. 4A). The parent-offspring regressions showed a significant parent–symbiont relationship in the lectins ConA (*P* = 0.023) and PNA (*P* = 0.028; Fig. 4B), while there was no significant effect in the other lectins (*P* > 0.05). For the significant lectins, R^2^ of the scaled parent-offspring regression were ConA (R^2^ = 0.415) and PNA (R^2^ = 0.027), indicating substantial explained variance. We observed near-zero or negative slopes for other lectins, which could reflect biological variation in glycomes across life history traits or may be representative of noise in a relatively small dataset. The slope of the parent–offspring regression was significantly different between symbiont genera in ConA (*P* = 0.038) and PNA (*P* = 0.032), but not other glycans (Fig. 4B). We observed several negative correlations between parent and egg MFIs in several combinations, but none were significant.

**Fig. 4.**
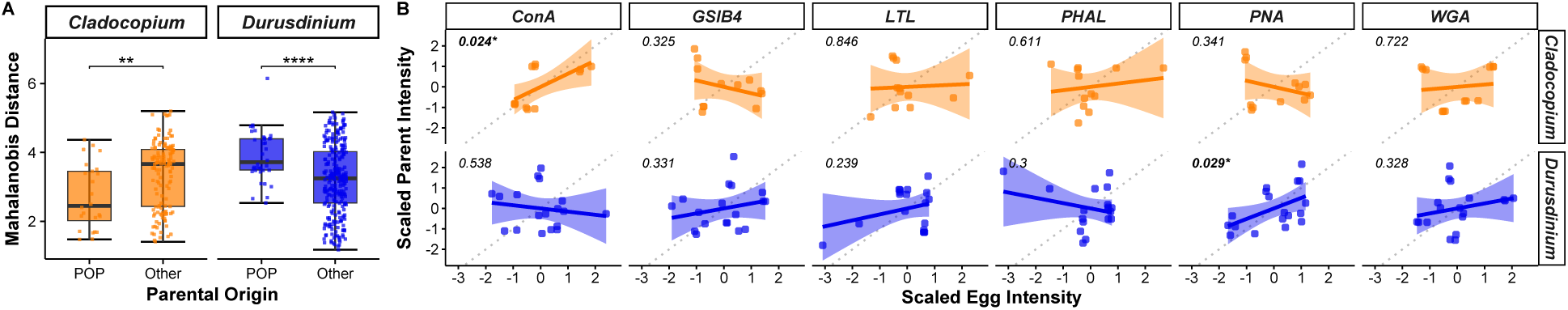
Lectin-binding profiles are conserved between symbiont generations. **A.** Mahalanobis distance distributions comparing lectin-binding profiles for symbiont cells associated with coral eggs. Distances were grouped by symbiont (*Cladocopium* and *Durusdinium*) and parental origin (parent-offspring population, POP; other populations, Other). Significance was assessed using Tukey-adjusted contrasts. **B.** Pairwise correlations between scaled lectin-binding intensities measured in symbionts isolated from coral eggs and their corresponding parental colonies, for *Cladocopium* (orange) and *Durusdinium* (blue). Each panel shows one lectin (ConA, GSIB4, LTL, PHAL, PNA, WGA). The dotted line represents the 1:1 identity line, and the solid line represents a linear regression fit with 95% confidence interval shading. Pearson correlation coefficients and significance are indicated. Asterisks denote adjusted *P* values (\**P*< 0.05, ** *P*< 0.01, *** *P*< 0.001).

## Discussion

The ability of coral reefs to persist under climate change depends in part on the composition and flexibility of corals’ symbiotic partnerships, which represent the energetic foundation of the ecosystem. Associations between Symbiodiniaceae and their cnidarian hosts rely on molecular specificity, which underpins the establishment and maintenance of the endosymbiosis. In vertically transmitting corals, symbionts are directly provisioned to offspring via eggs and are therefore expected to promote the continuity of symbiont communities across generations, increasing symbiont fidelity by coupling fitness between both partners (35). This method of symbiont transmission is hypothesized to arise from more mutually beneficial partnerships (36) and may reduce the likelihood of incompatible or marginal associations (37). However, it remains unclear whether the symbionts’ cellular traits are conserved during transmission and across host life stages, and whether they contribute to the stability of vertically transmitted symbioses.

Here, we examined cell-surface glycan profiles of *Cladocopium* and *Durusdinium* in parental colonies and eggs in the vertically transmitting coral *M. capitata*. The Symbiodiniaceae ITS2 sequences associated with eggs largely mirror those of their parental colonies in *Cladocopium*-and *Durusdinium*-dominated genotypes (10), indicating stable transmission at the community level. Our results showed that glycan signatures were structured by symbiont genus and differed between life stages, with mannose/glucose and galactose emerging as key contributors to this variation.

Egg-derived symbionts exhibited greater variability in lectin-binding profiles compared to those from parental colonies (Fig. 1C, D). This pattern was evident in both symbiont genera, but was more pronounced in *Durusdinium* (Fig. 1D). Increased variability in eggs suggests that glycan profiles may be less constrained at early life stages, potentially reflecting heterogeneity in symbiont populations prior to stabilization within the adult host environment. In contrast, the reduced variability observed in parental colonies may indicate more consistent or regulated glycan expression in established symbioses. This shift from higher variability in eggs to more uniform profiles in adults is consistent with increasing constraint in host–symbiont interactions over time (8) and provides a potential framework for interpreting the roles of specific glycans across life stages.

Mannose/glucose (ConA) and galactose (PNA) terminal glycans have been central to studies of Symbiodiniaceae cell–surface features and host–symbiont recognition. In our study, ConA exhibited the strongest MFI profile across samples (Fig. 1, 2; Supplementary Fig. 2), consistent with previous reports (14,20,24). This pattern likely reflects the abundance of mannose- and glucose-containing glycans on the Symbiodiniaceae surface (11,14,18,19), supported by the cellulose-rich dinoflagellate cell wall (21) and the enrichment of terminal mannose residues on surface glycoproteins (20). Functionally, mannose-rich glycans have been directly linked to symbiosis establishment and maintenance. Host-derived mannose-binding lectins localized around symbionts within the gastrodermis of *Acropora millepora* (38) and *Pocillopora damicornis* (39), and showed substantial variability in binding domains, suggesting a role in fine-scale recognition of compatible partners (38). Experimental saturation of *Exaiptasia diaphana* lectin receptors with D-mannose reduced colonization success by homologous symbionts (24), further supporting a role in recognition rather than a purely structural function in dinoflagellate cell walls.

D-galactose–containing glycans similarly contribute to symbiotic interactions. Galactose residues are present on the surface of multiple Symbiodiniaceae species (14,22,24,25) and have been implicated in symbiosis establishment in *A. tenuis* (40) larvae and anemone model (24). Two D-galactose-binding lectins, SLL-2 and CeCL, induced the transition of Symbiodiniaceae from a motile, flagellated state to a non-motile coccoid form characteristic of the symbiotic state, and localized around symbiont cells within host tissues (19,40–42). Together, these findings highlight both mannose/glucose- and galactose-containing glycans as key mediators of host–symbiont integration and emphasize the relative importance of galactose in gametes, where PNA is a stronger marker of symbiont variation and identity than in parents (Fig. 1, 4).

Multivariate differences between *Cladocopium* and *Durusdinium* were supported by pairwise correlations in lectin-binding intensities, with ConA and PNA – key drivers of genus-level separation – showing coordinated positive patterns in egg-derived symbionts (Fig. 3). In parental colonies, ConA and PNA correlations diverged between genera, positive in *Cladocopium*-dominated corals and negative in *Durusdinium*-dominated corals, although these relationships were not significant (Fig. 3). Previous work on *M. capitata* from Kāne‘ohe Bay showed that ConA binding profiles shift under thermal stress and are linked to oxidative stress, with similar trends observed for PNA (14). Considering oxidative stress can alter glycan composition through redox-mediated modification of glycoproteins (43), these observations suggest that symbiont cell-surface glycans may integrate physiological state with recognition processes.

Within this framework, our results provide evidence that the transmission of specific cell-surface glycans is conserved across life stages in *Cladocopium* and *Durusdinium*. The significant difference in Mahalanobis distances between matched parent–offspring pairs relative to unrelated comparisons (Fig. 4A) suggests that vertically transmitted *Cladocopium* symbionts retain genotype-specific glycan signatures, consistent with parental effects mediated through direct transmission of symbiont traits. However, this conservation is not uniform across the glycome. Lectin-level analyses revealed that only ConA and PNA exhibited significant parent–offspring correlations (Fig. 4B), and these were genus-specific, indicating that intergenerational similarity is driven by distinct glycan classes in each lineage. The high similarity in *Cladocopium*-dominated parents and eggs is likely driven by the strong heritability of mannose/glucose in this genus, which is also the most abundant glycan, weighting this analysis.

Notably, these same lectins were primary contributors to genus-level differences in eggs and parents (Fig. 1), linking divergence between symbiont genera with parent–offspring continuity. This suggests that some glycan features, primarily mannose/glucose (ConA specificity), that distinguish *Cladocopium* and *Durusdinium*, are also those most consistently maintained across life stages, and may therefore play a functional role in maintaining host–symbiont compatibility. Conversely, galactose and N-acetylglucosamine (PNA and WGA specificity, respectively) were disproportionately important for distinguishing symbionts in gametes (Fig. 1A), but these glycans are orthogonal to the axis of separation in adults (Fig. 1B), suggesting differential provisioning and function in early life histories. This type of variation across life history stages matches moderate heritability differences between symbiont genera, suggesting some degree of strategic provisioning that differs based on the type of symbiont and corresponds to the overall breakdown in correlation structure in the glycome of the two corals.

All pairwise correlations in *Cladocopium* adults and eggs were positive, although some were not significant, indicating that differentiation between individuals within this population is driven by variation in the intensity of the lectin-probe signal and presumably the relative abundance of the glycans on cell surfaces (Fig. 2A). Conversely, in *Durusdinium*, negative correlations suggest some tradeoffs in relative abundance (Fig. 2B), which may be indicative of stronger compositional variation marks in *Durusdinium*, especially in offspring where multiple strong negative correlations emerge that are not found in adults or in *Cladocopium*.

Together, our findings support a model in which vertical transmission preserves recognition-relevant glycan features while allowing flexibility in other components of the symbiont surface. Rather than broad inheritance of the glycome, specific glycan classes—likely those involved in host–lectin interactions—appear to be selectively maintained across generations. Future work should extend these analyses to larvae and juveniles to determine whether glycan traits are maintained at these life stages, and to test whether shifts in the ConA:PNA ratio—reflecting mannose/glucose vs. galactose glycans—mediate parental effects and the preferential association of *Durusdinium* under stress through glyco-redox–linked mechanisms. This selective retention may represent a mechanism by which corals ensure symbiont fidelity while retaining physiological plasticity under changing environmental conditions.

## Author Contributions

GT and CD conceived the study and analyzed data. GT, KH and CD secured funding. GT, NLF, ACV, SLR, IAA, EM, KS and CD performed the experiments. GT wrote the manuscript. All authors collected data and revised the manuscript.

## Acknowledgements

Hawaiian culture fosters a reciprocal connection between people and coral reefs, including those in Kāneʻohe Bay, O‘ahu, Hawai‘i where this research was conducted. We hope to honor this relationship by recognizing its foundational importance in our work. We are grateful to the Coral Resilience Lab for field and laboratory support. All collections were made under Hawaiʻi Department of Land and Natural Resources permits to HIMB (SAP 2026-16). We thank the Walder-Christensen Charitable Fund for supporting this work. This is SOEST contribution #xx and HIMB contribution #xx.

---

**Supplementary Fig. 1.**
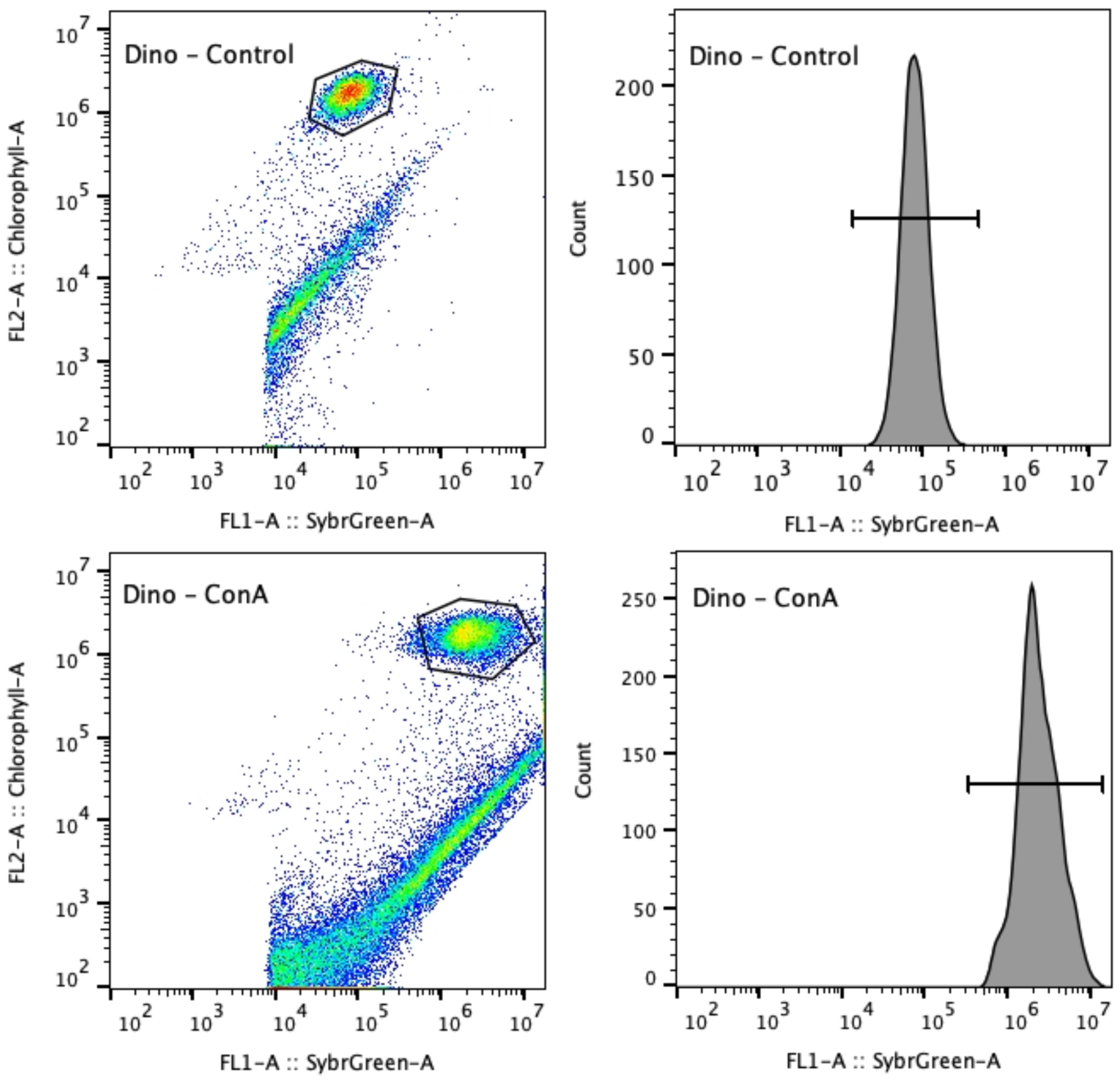
Flow cytometry gating and distribution of Symbiodiniaceae cells. Representative flow cytometry plots showing the identification and quantification of Symbiodiniaceae cells based on chlorophyll autofluorescence (top panel) and lectin molecular probe staining (bottom panel). Right panel shows density plots of FL2-A (Chlorophyll-A autofluorescence) versus FL1-A (SYBR Green fluorescence), with the gated population corresponding to Symbiodiniaceae cells for unstained cells (control) and stained cells (ConA). Colors indicate event density from low (blue) to high (red). Left panel shows histograms of FL1-A fluorescence intensity (SYBR Green) for the gated population, showing the distribution of lectin molecular probe signal. Axes are displayed on a log scale.

**Supplementary Fig. 2.**
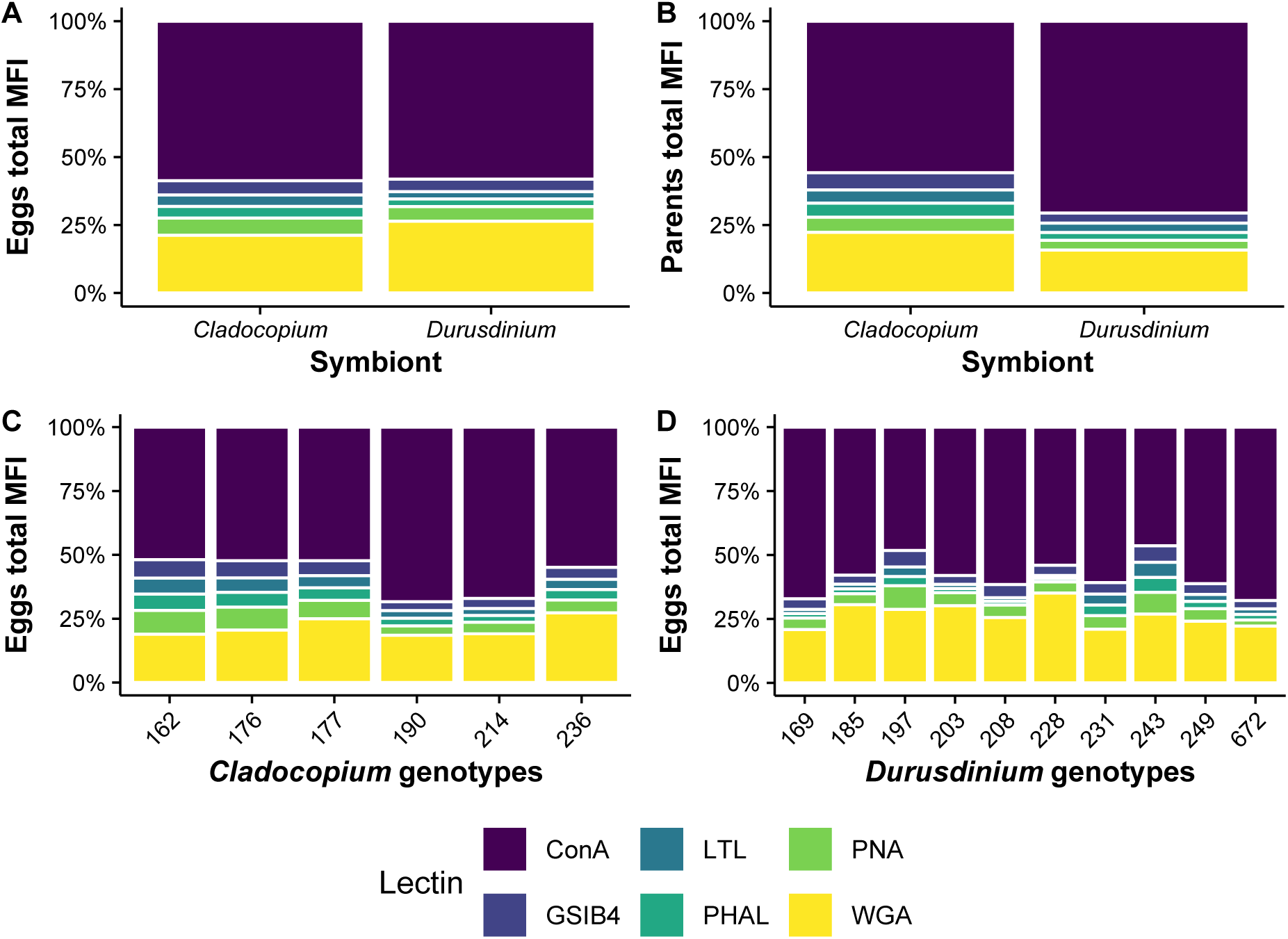
Median fluorescent intensity of Symbiodiniaceae cells. **A.** Proportional binding profiles (total positive MFI signal) across lectins in *Cladocopium* (*N* = 18) and *Durusdinium* (*N* = 30) isolated from coral eggs. **B.** Proportional binding profiles across lectins in *Cladocopium* (*N* = 15) and *Durusdinium* (*N* = 18) isolated from parental colonies. In C and D, bar height represents the relative contribution of each lectin to the total binding signal per symbiont genus. **C.** Proportional binding profiles across lectins per host genotype in *Cladocopium* (*N* = 18) isolated from coral eggs. **D.** Proportional binding profiles across lectins per host genotype in *Durusdinium* (*N* =30) isolated from coral eggs. In E and F, bar height represents the relative contribution of each lectin to the total binding signal per genotype. Lectins: ConA (concanavalin A, specific for D-mannose and D-glucose), LTL (*Lotus tetragonolobus* lectin, specific for L-fucose), PNA (*Arachis hypogaea* lectin, specific for D-galactose), WGA (wheat germ agglutinin, specific for N-acetylglucosamine and N-acetylneuraminic acid), PHA-L (phytohemagglutinin-L from *Phaseolus vulgaris*, specific for N-acetylglucosamine β(1-2) mannopyranosyl) and GS-IB4 (isolectin from *Griffonia simplicifolia*, specific for N-acetyl-D-galactosamine and a-D-galactosyl residues).

**Supplementary Table 1.**
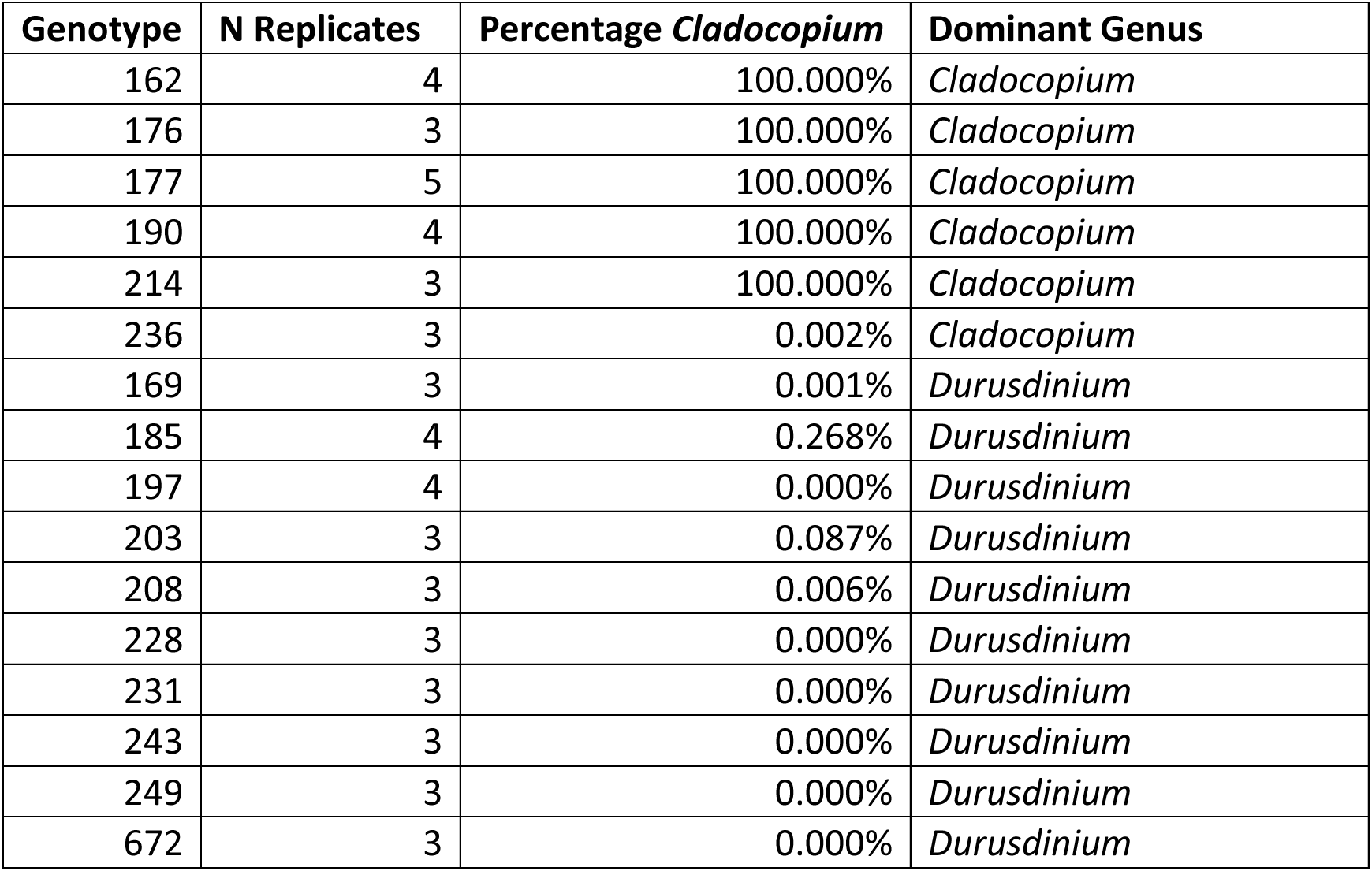
Symbiont densities in *M. capitata* colonies.

