## Supplementary material for "Selective conservation of symbiont cell-surface glycans across generations in a vertically transmitting coral"

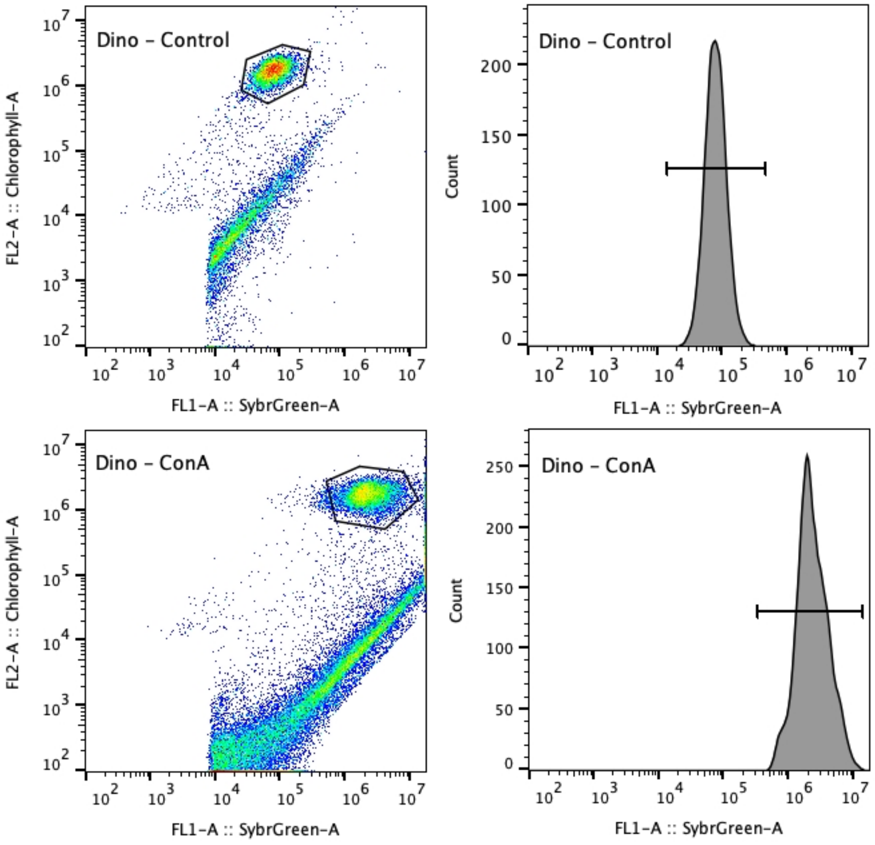


**Supplementary Fig. 1 | Flow cytometry gating and distribution of Symbiodiniaceae cells.**

Representative flow cytometry plots showing the identification and quantification of Symbiodiniaceae cells based on chlorophyll autofluorescence (top panel) and lectin molecular probe staining (bottom panel). Right panel shows density plots of FL2-A (Chlorophyll-A autofluorescence) versus FL1-A (SYBR Green fluorescence), with the gated population corresponding to Symbiodiniaceae cells for unstained cells (control) and stained cells (ConA). Colors indicate event density from low (blue) to high (red). Left panel shows histograms of FL1-A fluorescence intensity (SYBR Green) for the gated population, showing the distribution of lectin molecular probe signal. Axes are displayed on a log scale.

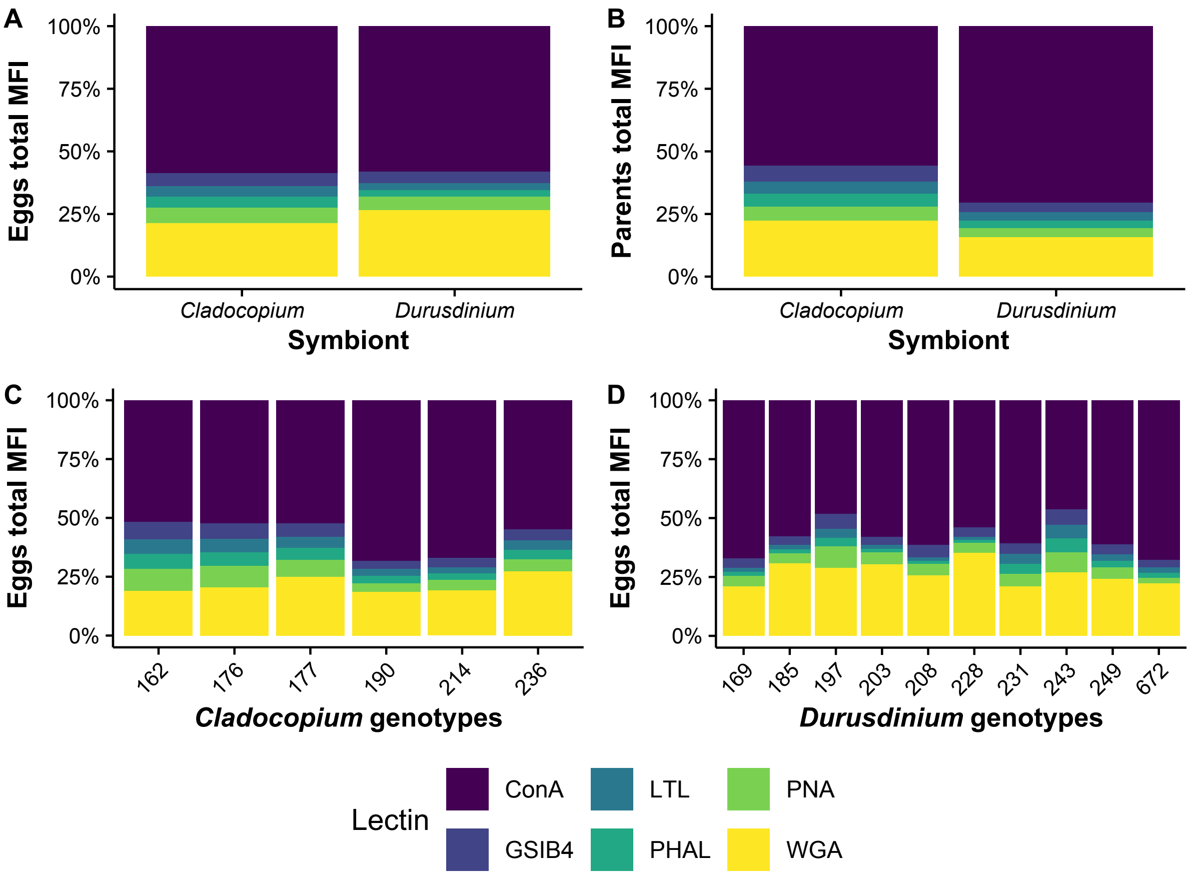


**Supplementary Fig. 2 | Median fluorescent intensity of Symbiodiniaceae cells.**

**A.** Proportional binding profiles (total positive MFI signal) across lectins in *Cladocopium* (*N* = 18) and *Durusdinium* (*N* = 30) isolated from coral eggs. **B.** Proportional binding profiles across lectins in *Cladocopium* (*N* = 15) and *Durusdinium* (*N* = 18) isolated from parental colonies. In C and D, bar height represents the relative contribution of each lectin to the total binding signal per symbiont genus. **C.** Proportional binding profiles across lectins per host genotype in *Cladocopium* (*N* = 18) isolated from coral eggs. **D.** Proportional binding profiles across lectins per host genotype in *Durusdinium* (*N* =30) isolated from coral eggs. In E and F, bar height represents the relative contribution of each lectin to the total binding signal per genotype. Lectins: ConA (concanavalin A, specific for D-mannose and D-glucose), LTL (*Lotus tetragonolobus* lectin, specific for L-fucose), PNA (*Arachis hypogaea* lectin, specific for D-galactose), WGA (wheat germ agglutinin, specific for N-acetylglucosamine and N-acetylneuraminic acid), PHA-L (phytohemagglutinin-L from *Phaseolus vulgaris*, specific for N-acetylglucosamine β(1-2) mannopyranosyl) and GS-IB4 (isolectin from *Griffonia simplicifolia*, specific for N-acetyl-D-galactosamine and a-D-galactosyl residues).

| Genotype | N Replicates | Percentage *Cladocopium* | Dominant Genus |
|---|---|---|---|
| 162 | 4 | 100.000% | *Cladocopium* |
| 176 | 3 | 100.000% | *Cladocopium* |
| 177 | 5 | 100.000% | *Cladocopium* |
| 190 | 4 | 100.000% | *Cladocopium* |
| 214 | 3 | 100.000% | *Cladocopium* |
| 236 | 3 | 0.002% | *Cladocopium* |
| 169 | 3 | 0.001% | *Durusdinium* |
| 185 | 4 | 0.268% | *Durusdinium* |
| 197 | 4 | 0.000% | *Durusdinium* |
| 203 | 3 | 0.087% | *Durusdinium* |
| 208 | 3 | 0.006% | *Durusdinium* |
| 228 | 3 | 0.000% | *Durusdinium* |
| 231 | 3 | 0.000% | *Durusdinium* |
| 243 | 3 | 0.000% | *Durusdinium* |
| 249 | 3 | 0.000% | *Durusdinium* |
| 672 | 3 | 0.000% | *Durusdinium* |

**Supplementary Table 1 |** Symbiont densities in *M. capitata* colonies.
